# Unraveling the Interplay of Temperature Adaptation, Lipidomics, and Environmental Factors in *Acinetobacter baumannii* Clinical Strains

**DOI:** 10.1101/2023.08.09.552612

**Authors:** Clara Dessenne, Benoît Ménart, Sébastien Acket, Gisèle Dewulf, Yann Guerardel, Olivier Vidal, Yannick Rossez

**Affiliations:** Université Lille, CNRS, UMR 8576-UGSF-Unité de Glycobiologie Structurale et Fonctionnelle, F-59000 Lille, France; Centre Hospitalier de valenciennes, Laboratoire de Biologie Hygiène - service de Microbiologie Avenue Désandrouins, 59300 Valenciennes; Université de technologie de Compiègne, UPJV, UMR CNRS 7025, Enzyme and Cell Engineering, Centre de recherche Royallieu -CS 60319 -60203 Compiègne Cedex, France; Institute for Glyco-core Research (iGCORE), Gifu University, Gifu, Japan

## Abstract

*Acinetobacter baumannii* has gained prominence due to its heightened antibiotic resistance and adaptability within healthcare settings. Unlike other *Acinetobacter* species, *A. baumannii* predominantly thrives within healthcare environments, where its persistence is underscored by physiological adaptations, including homeoviscous adaptation that modifies glycerophospholipids (GPL) to enhance membrane flexibility. The bacterium’s substantial genetic diversity highlights the paramount importance of prudent strain selection for research involving drug resistance and virulence. This study investigates the lipid composition of six clinical *A. baumannii* strains, incorporating the highly virulent model strain AB5075 with multiple antibiotic resistances. Our objective is to scrutinize the adaptations of glycerophospholipids (GPL) and glycerolipids (GL) within these isolated strains, each characterized by unique antibiotic resistance profiles, under variable temperature conditions mimicking environmental and physiological scenarios. The strains’ differential performance in motilities and biofilm formation across varying temperatures reveals intriguing patterns. Notably, the study uncovers a consistent elevation in palmitoleic acid (C16:1) content in five of the six strains at 18°C. Utilizing LC-HRMS^2^ analysis, we elucidate shifts in GPL and GL compositions as temperatures oscillate between 18°C and 37°C for all strains. Exploration of lipid subspecies further exposes disparities in PE and PG lipids containing C16:1 and oleic acid (C18:1). This investigation not only provides insights into the physiological attributes and survival strategies of *A. baumannii* but also deepens our comprehension of its adaptive responses to temperature changes. By unraveling the dynamics of lipid composition and fatty acid profiles, this study enriches our understanding of the bacterium’s ecological fitness and behavior in diverse environments.

**IMPORTANCE:** *Acinetobacter baumannii*, a bacterium known for its resistance to antibiotics, is a concern in healthcare settings. This study focused on understanding how this bacterium adapts to different temperatures and how its lipid composition changes. Lipids are like the building blocks of its cell membranes. By studying these changes, scientists can gain insights into how the bacterium survives and behaves in various environments. This knowledge helps us better understand its ability to cause infections and resist treatments. The study’s findings contribute to our broader understanding of how *Acinetobacter baumannii* functions, which is important for developing strategies to combat its impact on patient health.

## INTRODUCTION

*Acinetobacter* spp., a versatile and resilient group of bacteria, has gained significant attention in recent years due to mainly *Acinetobacter baumannii* which is commonly found in healthcare settings and long-term care facilities (1). This bacterium, a member of the *Acinetobacter calcoaceticus-baumannii* (Acb) complex, is implicated in human infections, alongside other species such as *A. lwoffi, A. junii, A. nosocomialis*, and *A. pittii*. Additionally, *A. calcoaceticus* is regarded as an environmental species (2). One of the biggest concerns with *A. baumannii* is its ability to develop resistance to multiple antibiotics, which can make infections difficult to treat (3). Although *Acinetobacter* spp. are renowned for their widespread presence and exceptional adaptability, the case appears to differ when it comes to *A. baumannii*. Unlike other *Acinetobacter* species that are commonly found in soil or water samples, A. *baumannii* is rarely isolated from such sources unless inadvertently introduced through human waste (4, 5). Instead, *A. baumannii* has demonstrated a higher propensity for survival within healthcare facilities (6, 7), potentially facilitated by transient colonization on the hands of healthcare workers and dissemination through aerosolized particles from infected patients (8, 9).

To thrive in hospital environments and survive outside the human body, *A. baumannii* undergoes physiological adaptations that can influence its pathogenicity, such as its ability to form biofilm or exhibit motility. Motility is a critical mechanism that aids *A. baumannii* in the dissemination within hospital environments. Additionally, motility also plays a significant role in actively moving towards and colonizing specific sites within the host (10). Two distinct types of motility have been characterized in *Acinetobacter* spp.: twitching motility, which relies on the type IV pilus for movement (11), and surface-associated motility. The latter is influenced by various factors, including the PrpABCD pilus and 1,3 diaminopropane (12–15). In addition to motility, *A. baumannii*’s capability to form biofilms further contributes to its pathogenicity and persistence by making eradication more challenging (16, 17). Interestingly, the impact of temperature on the physiology and virulence of *A. baumannii* remains relatively understudied while these nosocomial bacteria rely on a dynamic interplay between their ability to colonize the human body and their capacity to spread within the hospital environment. This intricate balance is crucial for their persistence and dissemination in healthcare settings. One prominent mechanism known as homeoviscous adaptation is a common response to low temperatures, enabling the bacteria to withstand extracorporeal conditions. This adaptation involves modifying the composition of glycerophospholipids (GPL) by altering their fatty acids, ultimately enhancing their flexibility (18). Therefore, the bacterial cell membrane plays a crucial role in providing a protective barrier against the external environment and importantly to antibiotic resistance in several bacterial species (19–21). GPL in *A. baumannii* are found through seven sub classes including phosphatidylethanolamine (PE), phosphatidylglycerol (PG), lysophosphatidylethanolamine (LPE), hemibismonoacylglycerophosphate (HBMP), cardiolipin (CL), monolysocardiolipin (MLCL) and phosphatidylcholine (PC) (22, 23). *A. baumannii* contains two glycerolipids (GL), namely triacylglycerol (TG) and diacylglycerol (DG) (22). These GL serve not only as reserve compounds but also fulfill roles in metabolism, potentially acting as sources of fatty acids that can be readily mobilized (24).

*A. baumannii*, shows high genetic diversity. Reference strains don’t represent the full range of clinical isolates, so strain selection is crucial. The use of contemporary isolates should be preferred over historical strains to ensure accurate research on virulence and drug resistance (25). In this article, five strains of *A. baumannii* isolated from patients in intensive care from France are compared with the highly virulent model strain, AB5075, isolated in 2008 from a combatant wound infection (26, 27). The primary goal of this research is to unravel the intricate relationship between temperature and the virulence-related characteristics of *A. baumannii*. Specifically, we seek to explore the influence of temperature on twitching motility and biofilm formation capabilities in six recently isolated clinical strains. In addition, we aim to comprehensively analyze the lipid composition of these strains, focusing on their fatty acid content and conducting liquid chromatography-high-resolution tandem mass spectrometry (LC-HRMS^2^) to delve deep into the GPL and GL profile. This comprehensive investigation will shed light on the lipidic adaptation of *A. baumannii* under temperature conditions that faithfully replicate those encountered within the human body and hospital environments. By uncovering the intricate interplay between temperature and *A. baumannii*’s pathogenic traits, this study will contribute to our understanding of the bacterial adaptation mechanisms and potentially provide valuable insights for the development of novel strategies to address this resilient pathogen in healthcare settings.

## RESULTS

### Bacterial species-level identification and antibiotic resistance pattern

Five strains of *Acinetobacter* spp. were isolated from France, specifically selected to represent different antibiotic resistance profiles. Additionally, a reference strain, AB5075, isolated from a patient with osteomyelitis in USA was included in this study (26). To identify the five isolated strains, matrix-assisted laser ionization time-of-flight mass spectrometry (MALDI-TOF MS) has been used. This method has been effectively utilized in clinical microbiology labs for rapid bacterial identification. However, this technique is unable to accurately identify species within the Acb complex (28). Although all strains were initially identified as *A. baumannii*, we conducted amplification and sequencing of the 16S-23S rRNA gene spacer region (29) to confirm the identity of the strains used in this study. Fragments of approximately 600 bp were observed, exhibiting a high degree of similarity among the six strains, including AB5075 (Supplementary figure 1). Furthermore, as indicated in table 1, the strain AB5075 exhibits resistance to nearly all antibiotics, with the exception of minocycline, as previously described (26). ABVal3 is resistant to all antibiotics tested whereas ABVal2 requires higher concentrations of Ticarcillin, Trimethoprim-sulfamethoxazole and three β-lactams (clavulanic acid with tricarcillin, cefepime and imipenem) to exhibit susceptibilities. ABVal1 is susceptible to Trimethoprim–sulfamethoxazole only while ABVal4 is resistant to Aztreonam, Fosfomycin and at low concentration to Ciprofloxacin.

**Table 1.** Antibiotic resistance exhibited by the strains used in this study. R indicates resistance to the corresponding antibiotic and S stands for susceptible.

| Antibiotic group | Antibiotics | ABVal1 | ABVal2 | ABVal3 | ABVal4 | ABVal5 | AB5075 |
| --- | --- | --- | --- | --- | --- | --- | --- |
| Carboxypenicillin | Ticarcillin | R | S (at high concentration) | R | S | R | R |
| Carboxypenicillin - β-lactam | Ticarcillin - clavulanic acid | R | S (at high concentration) | R | S | S | R |
| β-lactam | Piperacillin | R | R | R | S | R | R |
| β-lactam - β-lactamase inhibitor | Piperacillin - Tazobactam | R | R | R | S | R | R |
| β-lactam | Aztreonam | R | R | R | R | R | R |
| β-lactam | Ceftazidime | R | R | R | S | R | R |
| β-lactam | Cefepime | R | S (at high concentration) | R | S | R | R |
| β-lactam | Imipenem | R | S (at high concentration) | R | S | S | R |
| Fluoroquinolone | Levofloxacin | R | R | R | S | R | R |
| Fluoroquinolon | Ciprofloxacin | R | R | R | S (at high concentration) | R | R |
| Aminoglycoside | Gentamicin | R | R | R | S | R | R |
| Aminoglycoside | Tobramycin | R | R | R | S | R | R |
| Aminoglycoside | Amikacin | R | R | R | S | R | R |
| Tetracycline | Minocycline | R | R | R | S | S | S |
| Phosphonic | Fosfomicin | R | R | R | R | R | R |
| Antifolate - Sulfonamide | Trimethoprim - sulfamethoxazole | S | S (at high concentration) | R | S | R | R |

**Figure 1.**
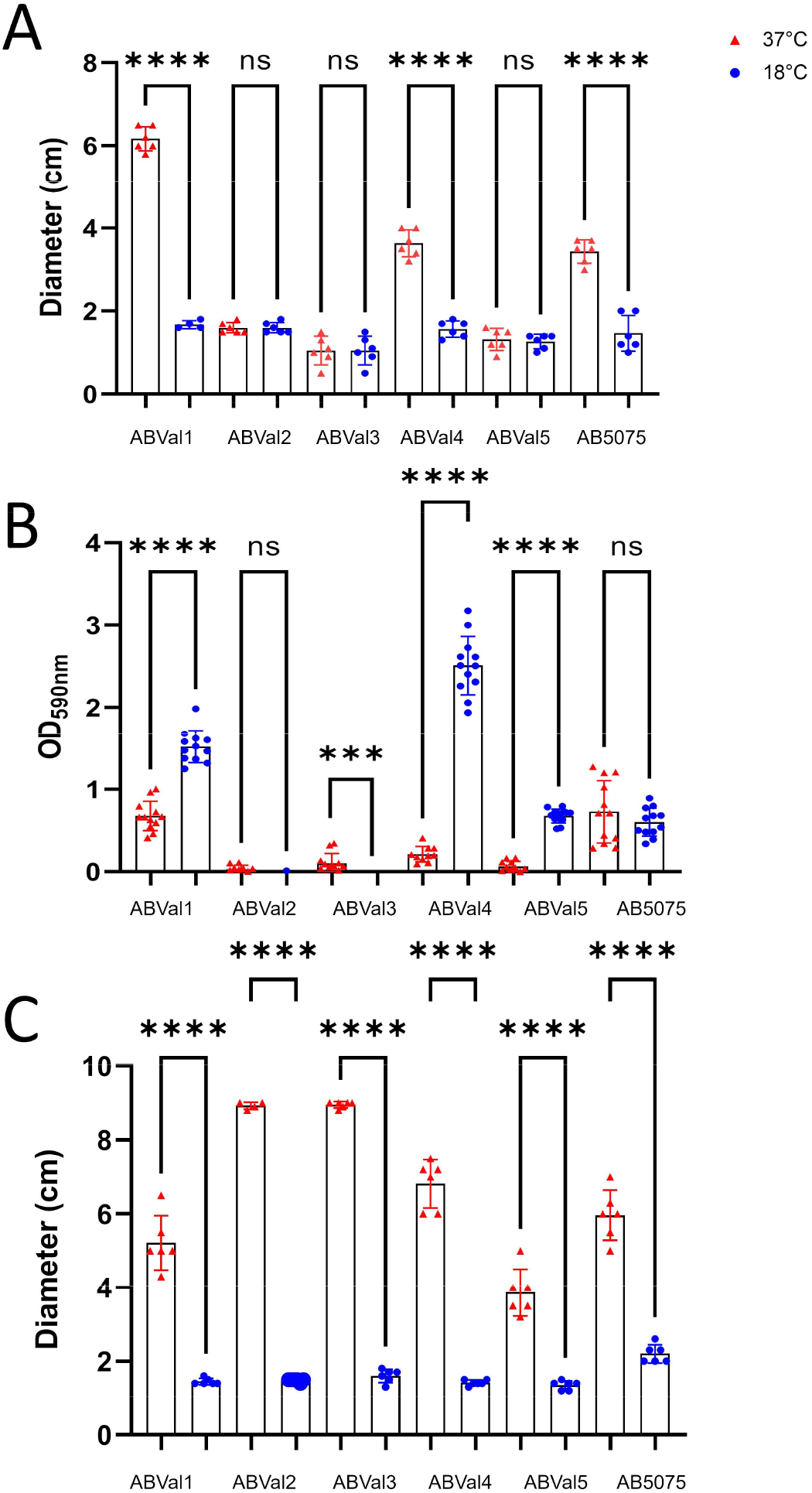
(A), Twitching motility at 37°C and 18°C after 48 hours of inoculation. (B), Biofilm formation at 37°C and 18°C. The graph represents the level of biofilm formation (absorbance at 590nm), which was investigated in a 96-well microtiter tray using a crystal violet stain method. (C), Surface associated motility at 37°C and 18°C after 48 hours of inoculation. The results correspond to at least 3 biologically independent samples. Statistical significances were determined by a two-tailed student’s *t* test (****, *p* ≤ 0.0001; ***, *p* ≤ 0.001; ns, *p* > 0.05).

### Motility and biofilm formation

To comprehensively assess bacterial behavior at both 37°C and 18°C, we evaluated the twitching motility of all six strains (Figure 1A). Among the strains, ABVal2, ABVal3, and ABVal5 showed minimal motility at either temperature, whereas ABVal1, ABVal4, and AB5075 exhibited significant twitching motility at 37°C. Of these, ABVal1 displayed particularly efficient movement, covering a greater distance than the others at 37°C. Notably, no distinct twitching motility was observed at 18°C.

Given that Type IV pili have been proposed to play a role in biofilm formation by facilitating initial bacterial attachment, a concept demonstrated in various bacterial species, including *A. baumannii* (11, 30, 31), we proceeded to evaluate each strain’s biofilm-forming capacity at both temperatures (Figure 1B). ABVal2 and ABVal3 showed negligible biofilm formation at 18°C and modest biofilm production at 37°C, with values slightly exceeding 0.28 OD590. Interestingly, AB5075 exhibited consistent medium-level biofilm production that remained unaffected by the temperature variations. ABVal5 displayed medium biofilm production, but exclusively at 18°C, while its biofilm formation weakened significantly at 37°C. ABVal1 and ABVal4 emerged as robust biofilm producers at 18°C; however, their biofilm production reduced to a medium level at 37°C. As described before, there appeared to be an inverse correlation between surface-associated motility and the capacity to form biofilms (32). At 18°C, motility was virtually absent across all strains, while at the same time, motility was evident in all strains (Figure 1C). Notably, ABVal2 and ABVal3 demonstrated the highest motility among the strains.

### Fatty acids content at different temperatures

To better estimate the temperature adaptation of the six strains, the fatty acid content was determined using gas chromatography with a flame ionization detector (GC-FID) after culturing them in LB at 37°C and 18°C (Figure 2). At 37°C, the major fatty acids detected were palmitic acid (C16:0) and oleic acid (C18:1), comprising approximately 35-40% and 30-40% of the total fatty acids, respectively. ABVal2, however, exhibited approximately 30% and 10% of these two fatty acids. Interestingly, ABVal2 also displayed a higher percentage of palmitoleic acid (C16:1) compared to other strains, with levels around 40% (Figure 2B). In contrast, C16:1 content varied between 5% and 15% at 37°C in the other strains, while the remaining fatty acids were below 5%, consistent with previous observations (22). At 18°C, a significant increase in C16:1 content was observed in strains ABVal1, ABVal3 to ABVal5, and AB5075 compared to 37°C (Figure A, C, E and F respectively). Notably, ABVal2 exhibited a substantial increase in oleic acid content at 18°C compared to 37°C. In order to achieve a more comprehensive comprehension of the apparent differences at 37°C and 18°C, particularly concerning ABVal2, we delved into the GPL and GL composition beyond just fatty acid content, employing LC-HRMS2 analysis.

**Figure 2.**
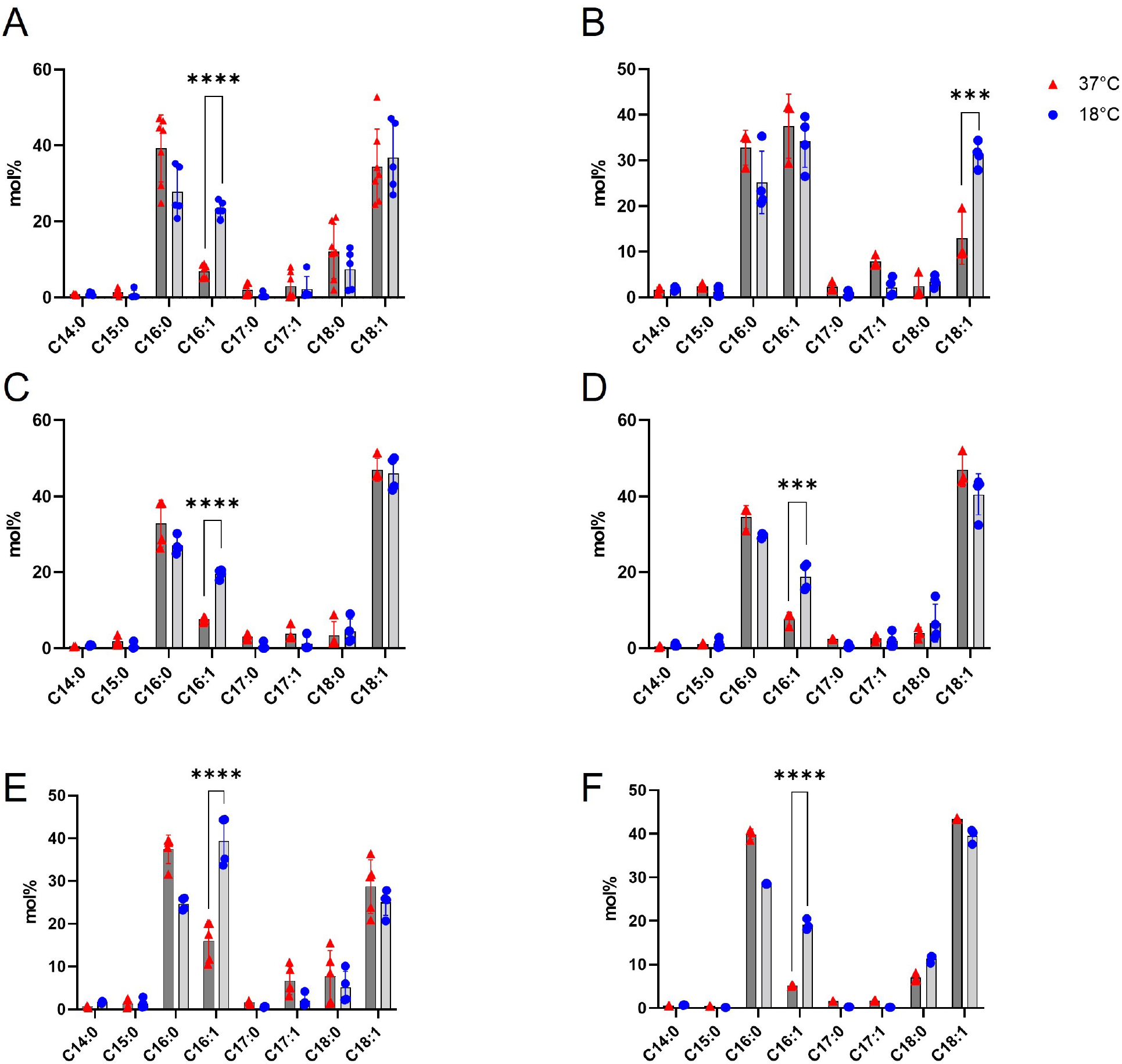
Fatty acid methyl ester analysis of *A. baumannii* clinical strains by GC-FID cultivated at 37°C or 18°C. The results correspond to n = 3 biologically independent samples. (A), ABVal1; (B), ABVal2; (C), ABVal;3 (D), ABVal4; (E), ABVal5; (F), AB5075. Statistical significances were determined by a two-tailed student’s *t* test (****, *p* ≤ 0.0001; ***, *p* ≤ 0.001).

### LC-HRMS^2^ analyses and temperature adaptations

Partial least squares-discriminant analysis (PLS-DA) was utilized to analyze the lipidome data and determine if there was any discernible separation among each strain when exposed to 37°C and 18°C (Figure 3). While there were variations in the GPL and GL compositions between the two temperatures for all strains, ABVal1, ABVal2, and AB5075 displayed more pronounced differences (Figures 3A, B and F respectively).

**Figure 3.**
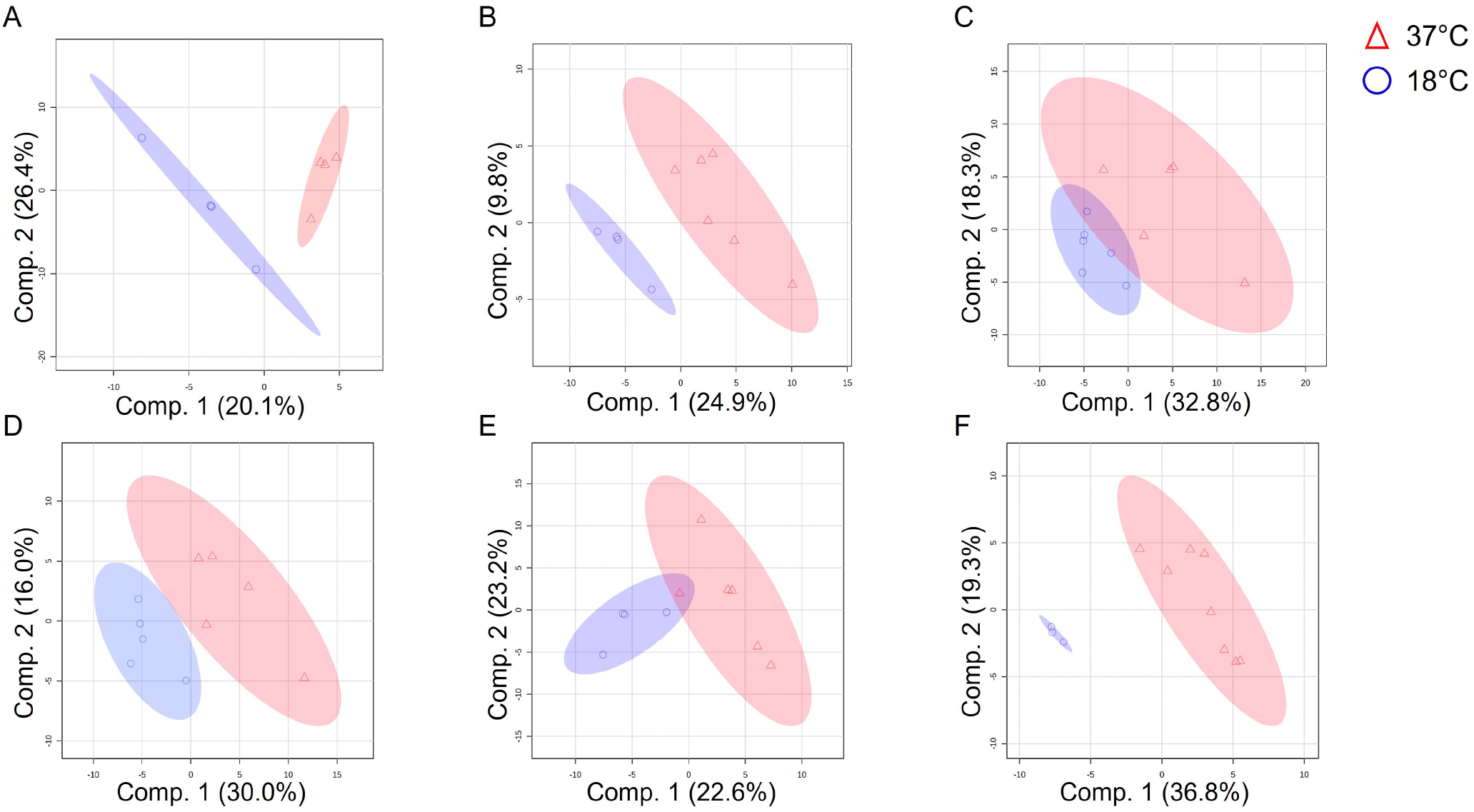
Partial least squares-discriminant analysis (PLS-DA) scores plot showing variances in lipid species between 37°C and 18°C for the strains (A), ABVal1; (B), ABVal2; (C), ABVal3; (D), ABVal4; (E), ABVal5; (F), AB5075. The results correspond to at least 3 biologically independent samples. The analysis was performed using MetaboAnalyst V5.0 (https://www.metaboanalyst.ca/, accessed in May 2023).

To identify the most upregulated lipids at 18°C for each strain, the data were analyzed using Volcano plots. The significance of the lipid with the highest fold-change was calculated, and these values were directly added to the scatterplot for clarity (Figure 4). For ABVal1 5 lipids were found to be upregulated at 18°C containing one or two C16:1 as follow: PG 16:1_18:1, CL 16:1_18:1_16:1_18:1, PG 16:1_16:1, HBMP 16:1/16:1_18:1 and PE 16:1_16:1. The first four are significantly different from 37°C (Figure 4A). ABVal2 showed only one lipid upregulated at 18°C, PE 16:1_18:1 (Figure 4B). While ABVal3 presented 3 lipids: PE 16:1_18:1, PG 16:1_18:1 and PE 16:1_16:1 with only the first significantly upregulated (Figure 4C). ABVal4 showed 4 lipids PE 16:1_16:1, PG 16:1_18:1, PG 16:1_16:1 and PA 16:0_16:1 with the last one non-significant (Figure 4D). ABVal5 and AB5075 had more upregulated lipids at 18°C with respectively 10 and 8 species. ABVal5 had HBMP 16:1/16:1_16:1, MLCL 16:1/16:1_16:1, PG 14:0_16:1, PE 14:0_16:1, PG 16:1_16:1, HBMP 16:1/16:1_18:1, CL 16:1_18:1_16:1_18:1, PE 16:1_16:1, CL 16:1_16:1_16:1_16:1 and MLCL 16:0_30:2. Among them only the two last lipids were not significant (Figure 4E). For AB5075 all the following lipids were significantly up regulated at 18°C: PG 16:0_16:1, HBMP 18:1/16:1_18:1, PA 16:1_16:1, PA 16:1_18:1, PC 18:1_18:1, PC 34:2, PG 16:1_18:1 and PE 16:1_16:1 (Figure 4F).

**Figure 4.**
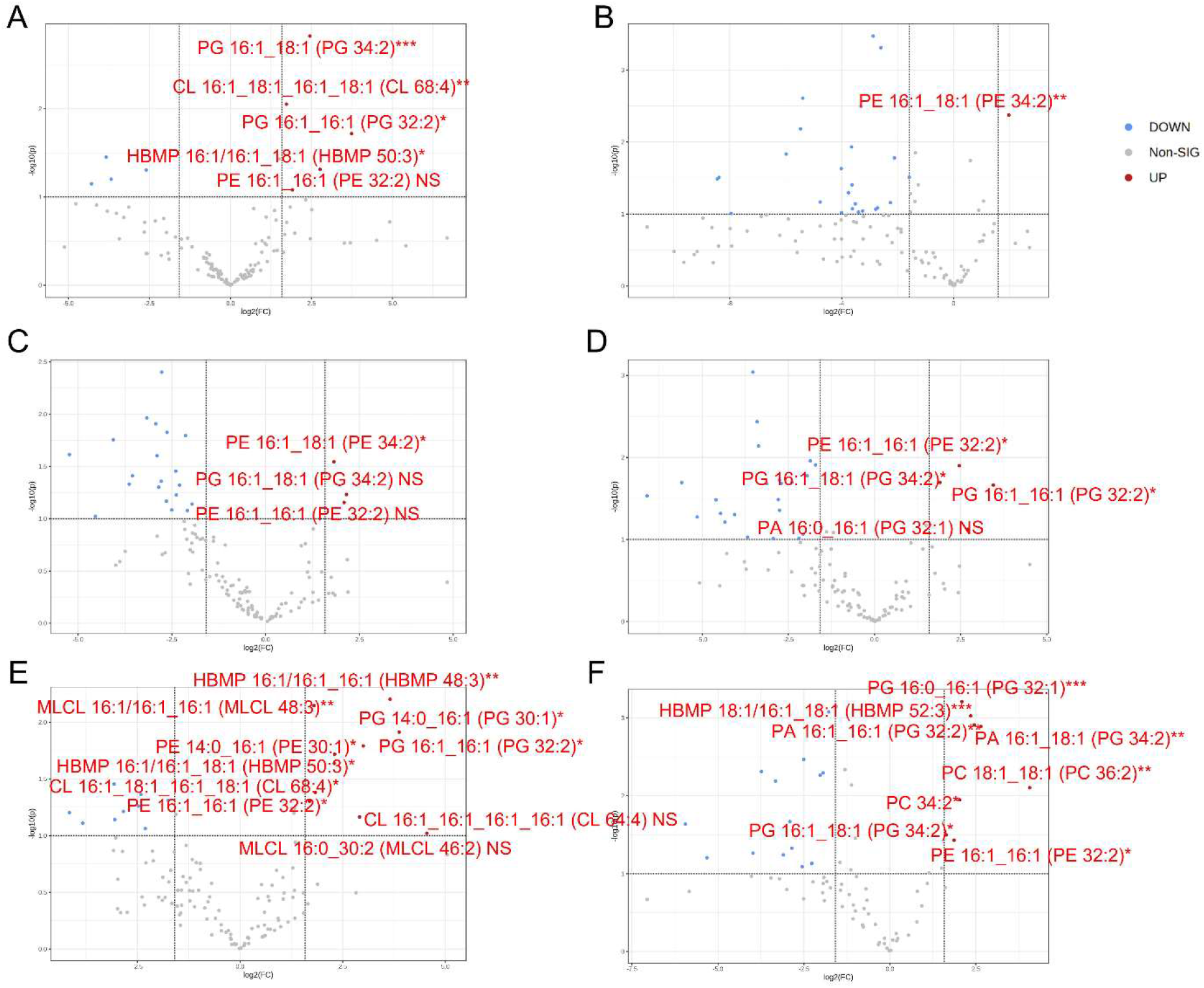
Volcano plot depicting lipids diversity of each strain at 37°C and 18°C with (A), ABVal1; (B), ABVal2; (C), ABVal;3 (D), ABVal4; (E), ABVal5; (F), AB5075. In red, all the lipids upregulated at 18°C are shown, along with their significance levels. The results correspond to at least 3 biologically independent samples. The analysis was performed using MetaboAnalyst V5.0 (https://www.metaboanalyst.ca/, accessed in May 2023). Statistical significances were determined by a two-tailed student’s *t* test (***, *p* ≤ 0.001; **, *p* ≤ 0.01; *, *p* ≤ 0.05 ; ns, *p* > 0.05).

Subsequently, to obtain a holistic understanding of each subclass, their respective proportions were computed (Figure 5). Across all strains, the prevailing lipids were predominantly PE and PG. Notably, there was a substantial rise in PE levels at 18°C for ABVal1, ABVal2, and ABVal3 (Figure 5A, B, and C, respectively). Conversely, in the case of AB5075, a reduction in PE content was discernible at 18°C, accompanied by a concurrent elevation in PG levels (Figure 5F).

**Figure 5.**
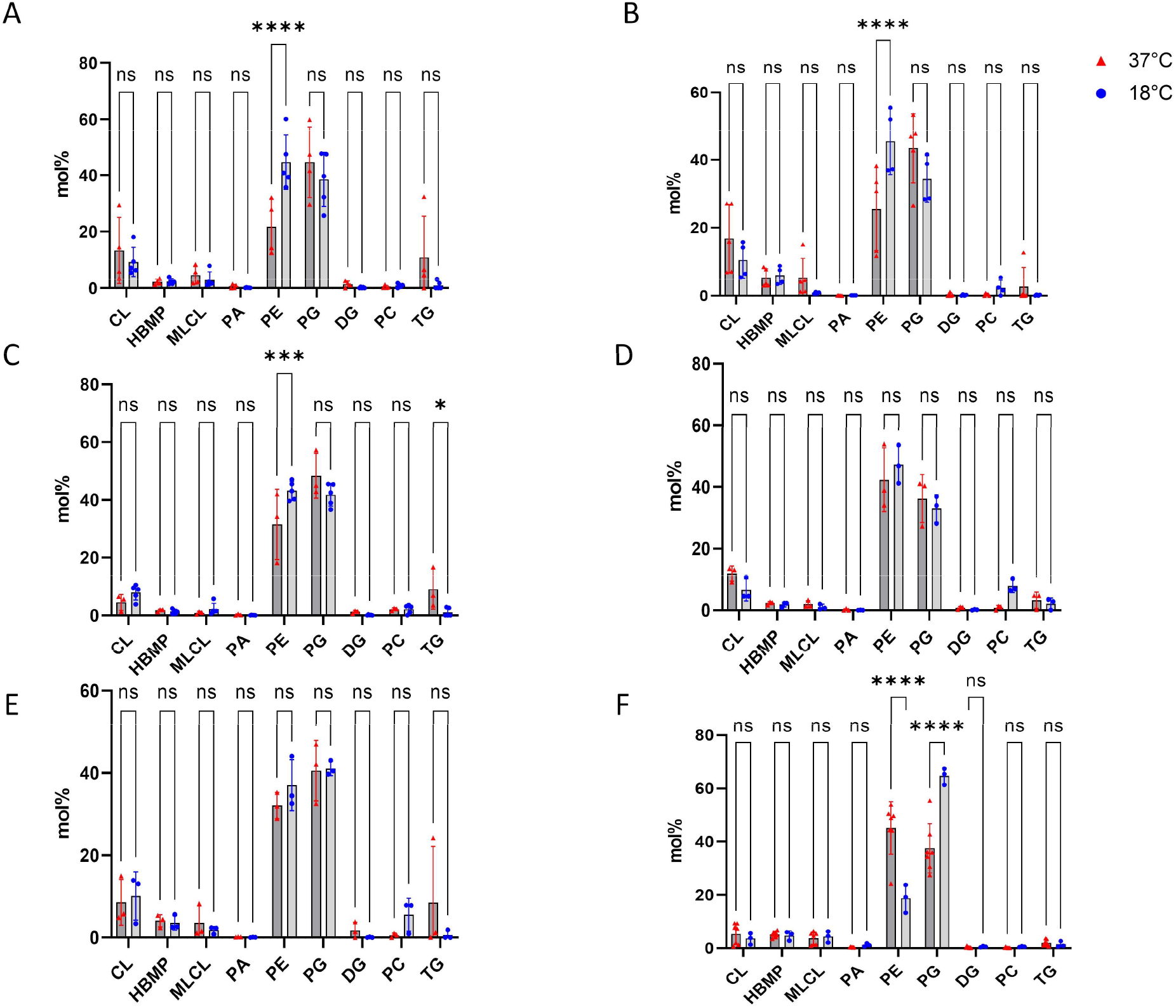
GPL and GL percentages of *A. baumannii* clinical strains cultivated at 37°C or 18°C based on LC-HRMS^2^ data with (A), ABVal1; (B), ABVal2; (C), ABVal;3 (D), ABVal4; (E), ABVal5; (F), AB5075. The results correspond to at least 3 biologically independent samples. Statistical significances were determined by *post-hoc* Tukey test after two-ways analysis of variance (****, *p* ≤ 0.0001; ***, *p* ≤ 0.001; ns, *p* > 0.05).

Given the prevailing abundance of PE and PG among the extracted lipids, the proportions of PE and PG molecules each incorporating one or two C18:1 and C16:1 fatty acids, which experience upregulation at 18°C, were computed and subsequently juxtaposed (Figure 6). In the case of ABVal1, 2, 3, 4, and 5, an elevated occurrence of PE containing C18:1 was discernible at 18°C, while no significant differences were apparent for PG (depicted by the white bars in figure 6A, B, C, D, and E, respectively). Conversely, for AB5075, the quantity of PE containing C18:1 exhibited a reduction, and although no significant effect was observed on PG, a similar discernible trend was distinctly evident, paralleling the pattern witnessed with PE (white bars in Figure 6F). The identical observation was repeated for PE containing C16:1 (represented by the grey bars in Figure 6A, B, C, D, E, and F, respectively). However, for PG containing C16:1, an elevation was noticed in ABVal1, 3, 4, 5, and AB5075. Particularly noteworthy was the more pronounced upsurge of PG C16:1 detected at 18°C for AB5075 (as indicated by the grey bars in Figure 6F). Intriguingly, for ABVal2, no significant fluctuation in either PE or PG containing C16:1 was discernible between the two temperatures (Figure 6B).

**Figure 6.**
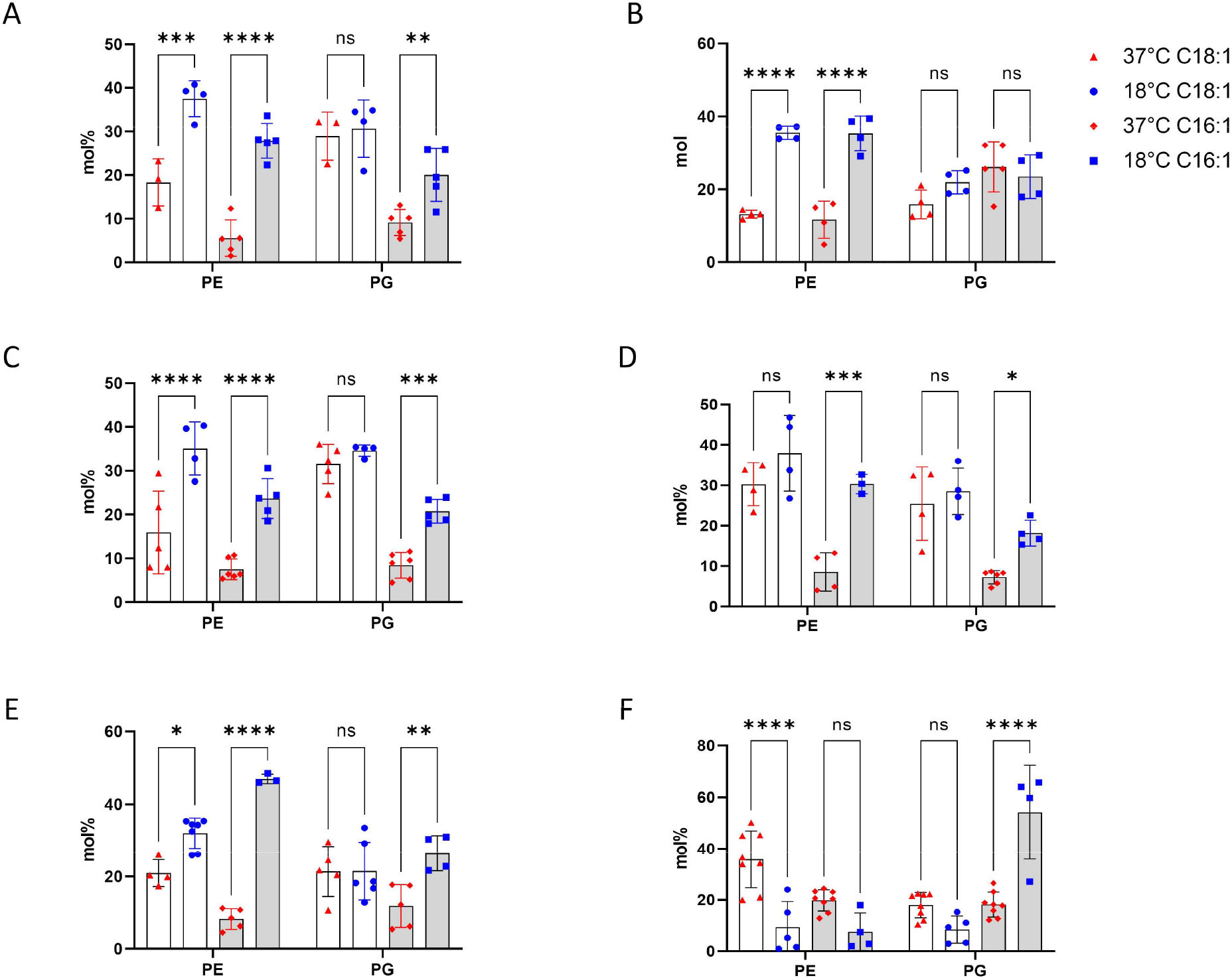
PE and PG containing at least one C16:1 or C18:1 percentages of *A. baumannii* clinical strains cultivated at 37°C or 18°C based on LC-HRMS^2^ data with (A), ABVal1; (B), ABVal2; (C), ABVal;3 (D), ABVal4; (E), ABVal5; (F), AB5075. The results correspond to at least 3 biologically independent samples. Statistical significances were determined by *post-hoc* Tukey test after two-ways analysis of variance (****, *p* ≤ 0.0001; ***, *p* ≤ 0.001; **, *p* ≤0.01; *, *p* ≤0.05; ns, *p* > 0.05).

## DISCUSSION

Like other bacterial species, *A. baumannii* maintains its protective barrier’s integrity through well-coordinated lipid homeostasis pathways, resulting in a diverse array of GPL and GL molecules (33). These major GPLs not only preserve membrane integrity but also aid in forming domains that facilitate protein translocation, guide cell division locations, and influence antibiotic penetration (22, 34–37). Therefore, understanding how lipids are formed in different *A. baumannii* strains with varying antibiotic susceptibility and how their synthesis is regulated under diverse physiological conditions is highly significant. In this study, our aim was to compare the physiological adaptations to GPL and GL compositions in six clinically isolated strains, each characterized by unique antibiotic resistance profiles (Table 1), under conditions mimicking environmental and body temperatures. The study reveals variations in twitching motility and biofilm formation capacities among the six strains at different temperatures (Figure 1). At lower temperatures, none of the strains exhibit motility, while three of them display enhanced biofilm-forming capabilities. ABVal1 showcases superior twitching motility at 37°C, with ABVal4 and AB5075 showing similar albeit lesser levels. Concerning biofilm formation, ABVal2 and ABVal3 exhibit minimal production, AB5075 consistently produces a medium amount, ABVal5’s production varies with temperature, and ABVal1 and ABVal4 display medium production at 37°C. The *Acb* complex has been associated with higher infection rates during warmer periods, as noted in prior studies (38). This observation aligns with the fact that *A. baumannii* infection incidence among hospitalized patients exhibits a peak during warmer months (39). This trend might be partly explained by the temperature-sensitive regulation of motility, as evidenced by its reduced efficiency when subjected to lower temperatures. Similarly, when examining the ATCC strain 17978, which was isolated in 1951 (40), research has uncovered that it exhibits an increased ability for biofilm formation at 28°C, accompanied by a decrease in its twitching motility. As previously mentioned, there exists a reverse relationship between surface-associated motility and biofilm formation, a point reiterated as well here (32, 41, 42). Globally, the specific motility and biofilm formation behaviors unique to each strain among the examined *A. baumannii* strains have been previously emphasized in studies involving various clinal strains (43–45).

The range of documented lipid variations in *A. baumannii* upon exposure to lower temperatures is limited. As far as our knowledge extends, modifications in fatty acid content under cold conditions have only been described in lipooligosaccharides (46). The present work sheds light on the matter by uncovering a significant increase in palmitoleic acid (C16:1) content for ABVal1, ABVal3-ABVal5, and AB5075 at 18°C. Whereas, ABVal2 exhibited a notable elevation in oleic acid (C18:1) content at 18°C (Figure 2). These findings imply that ABVal2 could exhibits a distinct desaturase specificity. Desaturases are enzymes that use oxygen to introduce double bonds into saturated fatty acyl chains, and their activity relies on an electron transport chain (47). However, in many bacteria, fatty acid desaturations can occur differently. Instead of specific desaturases, a double bond is introduced into a ten-carbon intermediate through the action of the fabA gene-encoded enzyme, which is part of the bacterial type II fatty acid biosynthetic pathway. This intermediate is then elongated to form 16:1 and 18:1 fatty acids (48–50). A recent study revealed poor desaturase activity in *Pseudomonas putida*, likely due to a weakly active component involved in the electron transfer process (51). Similarly, in ABVal2, the desaturase DesA identified in *A. baumannii*, which is characterized as desaturating C16:0 (52), could potentially fail to produce C16:1 and therefore be replaced by the activity of FabA, leading to the production of C18:1.

In the spectrum of tested antibiotics, ABVal4 displayed the least resistance compared to the other strains. Notably, this strain exhibited a remarkable proficiency in twitching motility, particularly conspicuous at 37°C, while it displayed its most optimal biofilm formation potential at 18°C. It’s worth noting that prior studies have established a link between heightened antibiotic resistance and increased biofilm formation by compiling data on more than 100 strains (53). However, drawing a definite conclusion solely from one strain is excessively speculative; a more comprehensive understanding necessitates the inclusion of additional strains to ensure statistically significant results.

Subsequently, LC-HRMS2 analyses were conducted to pinpoint the specific lipids that could potentially accommodate the heightened presence of unsaturated fatty acids at 18°C. Initially, a comprehensive assessment of the overall content was undertaken using PLS-DA, with the aim of discerning potential distinctions among various strains exposed to temperatures of 37°C and 18°C (Figure 3). The outcomes highlighted notable shifts in both GPL and GL compositions between the two temperatures for all strains. This observation aligns well with existing literature on bacterial adaptation facilitated by alterations in their lipid content (18, 54). Contrasting with a prior investigation, LPE displayed irregular detection across biological replicates, resulting in its omission from the analysis. In contrast, although phosphatidic acid (PA) was identified and included in this study, it comprised a mere fraction of the total lipid content though (22) (Figure 5).

Subsequently, we undertook an endeavor to identify lipid species that demonstrate an increase in expression at 18°C. However, among the outcomes, a consistent upregulation of lipid species across all strains at 18°C was not evident (Figure 4). As a result, we initiated an analysis at the lipid subspecies level, initially identifying the most prevalent lipids as a percentage across the strains and tested conditions (Figure 5). Subsequently, we concentrated on the two principal unsaturated fatty acids, C18:1 and C16:1 (Figure 2), which were upregulated at 18°C, specifically within the PE and PG lipids, known for their abundant presence (Figure 6). This analytical approach enabled us to deduce which lipids subspecies were accountable for the distinct fatty acid profile observed in ABVal2 (Figure 2). While PE containing C18:1 and C16:1 experienced upregulation at 18°C, no corresponding increase in PE and PG with augmented C16:1 content was noted. This contrasted with all other strains, which demonstrated elevated levels of PE and PG containing C16:1. Of note, AB5075 demonstrates a parallel elevation in C16:1 at 18°C (Figure 2F), with the exception of ABVal2. This increase is primarily attributed to PG at 18°C. Additionally, the diminishing levels of PE C18:1, PE C16:1, and PG C18:1 at 18°C indicate a discernible difference in lipid homeostasis compared to the other five strains under investigation. Hence, within the tested set of six strains, there are distinct behaviors observed: two strains exhibit dissimilar patterns, while the remaining four strains demonstrate notable similarity in terms of lipid synthesis under low temperatures. This phenomenon is a result of the significant gene repertoire diversity in *A. baumannii* and the notable influence of natural selection on protein evolution, driven by recombination and lateral gene transfer events within *A. baumannii* strains (55–58).

Overall, these findings emphasize the importance of temperature in influencing the lipid composition and fatty acid profiles of the bacterial strains studied. The observed variations highlight strain-specific differences as well as temperature-dependent adaptations, which could potentially have implications for the strains’ physiological properties, growth, and survival strategies. Further investigation into the functional implications of these lipidomic variations may provide insights into the mechanisms underlying the bacterial response to temperature changes and their overall ecological fitness.

## MATERIALS AND METHODS

### Strains and growth conditions

Clinical isolates were obtained from the microbiology department of the hospital of Valenciennes (France). All strains were isolated from the intensive care unit between 2014-2020. They were cultivated in lysogeny broth (LB) at 18°C or 37°C at 180 rpm. 18°C was considered as a relevant environmental temperature.

### Motility assays

All LB agar plates (1% concentration for twitching motility and 0.3% for surface-associated motility with Eiken chemical, Japan) were prepared freshly prior to being inoculated with 5 μL of a bacterial suspension in the exponential phase of growth (OD600: 0.3). For each temperature condition, the bacteria were initially cultured overnight at either 37°C or 18°C accordingly. The plates were then incubated at their respective temperatures for a duration of 48 hours. To ensure robustness of the results, three independent measurements were conducted for each strain.

### Biofilm assays

Prior to initiating the experiments, the bacteria were cultured overnight at either 37°C or 18°C. Subsequently, 100 μL of LB medium containing bacteria in the exponential growth phase were aseptically transferred to individual wells of a 96-well plate. The methodology used here is based on previous work (59), with slight modifications. The adherent cells underwent a series of steps. Initially, they underwent three washes with phosphate-buffered saline (PBS) and were then left to air-dry in a cabinet for two hours. Following this, the cells were stained by incubating them with a solution of 0.1% crystal violet for 10 minutes. After staining, the wells were washed three times with PBS to eliminate any excess dye. For dye release from the biofilm, a solution of ethanol containing 10% acetone was employed. The released dye’s absorbance was measured at 590nm using a plate reader. The presented biofilm data is an average taken from 10 or 12 wells. To assess the strains’ biofilm-forming ability, a modified version of a method derived from previous research (53) was employed. The average optical density at 590nm (OD_590nm_) of 48 control wells incubated without bacteria at both 37°C and 18°C was determined as 0.28 (ODc). Using this control value, the bacteria’s biofilm-forming capability was classified as strong when the value was 4 times ODc, medium between 2 times ODc and 4 times ODc, low if below 2 times ODc, and absence of biofilm if the values were equal to or below ODc.

### Antimicrobial susceptibility testing

The disk diffusion method was performed according to norms of Clinical Laboratory Standards Institute (CSLI) (60). Antibiotics tested included Ticarcillin (75 μg), Ticarcillin -clavulanic acid 7.5:1 (85 μg), Piperacillin (100 μg), Piperacillin – Tazobactam 1:1 (110 μg), Aztreonam (30 μg), Ceftazidime (30 μg), Cefepime (30 μg), Imipenem (10 μg), Levofloxacin (5 μg), Ciprofloxacin (5 μg), Gentamicin (10 μg), Tobramycin (10 μg), Amikacin (30 μg), Minocycline (30 μg), Fosfomycin (200 μg), Trimethoprim – sulfamethoxazole 1:19 (25 μg)

### MALDI-TOF identification

After an incubation of 72 h at 37 °C, single colonies were observed, and the bacterial species were identified by matrix-assisted laser desorption ionization-time-of-flight mass spectrometry (MALDI-TOF MS). Fresh colony material was spread on a MALDI target plate (MSP 96 target polished steel BC) (Bruker Daltonik GmbH, Germany) using a toothpick, mixed with 1 μL of a saturated α-cyano-4-hydroxy-cinnamic acid matrix solution in acetonitrile 50%–trifluoroacetic acid 2.5%, and dried in air at ambient temperature. Mass spectra were acquired and analyzed on a microflex LT/SH mass spectrometer (Bruker Daltonik) using a Bruker’s MALDI Biotyper software reference database library of 3995 entries, version 3.1.2.0 and default parameter settings, as reported.

### Amplification of ribosomal DNA

Amplification of 16S rRNA and 23S rRNA was performed with the specific universal primers 1512F (5GTCGTAACAAGGTAGCCGTA 3) and 6R (5GGGTTYCCCCRTTCRGAAAT3) as described previously (29).

### Lipid extraction

Lipids were extracted using the Bligh and Dyer method with a mixture of methanol (MeOH), chloroform (CHCl3), and water (H_2_O) in a volumetric ratio of 2:1:0.8 as described previously (22). The extracted lipids were stored at −20 °C until further analyses.

### Fatty acids analysis

The purified lipids were suspended in chloroform and a mixture of H2SO4 in methanol, butylated hydroxytoluene (BHT), and toluene was added and heated. Fatty acid methyl esters (FAME) were extracted using sodium chloride and heptane. The FAME composition was determined using gas chromatography, on a BPX70 column as described previously (22).

### LC-HRMS^2^, data processing and annotation

The liquid chromatography used a Waters Aquity UPLC C18 column (100 × 2.4 mm, 1.7 μm) coupled to an Acquity UPLC CSH C18 VanGuard precolumn (5 × 2.1 mm; 1.7 μm) at 65 °C. The mobile phase 60:40 (vol/vol) acetonitrile/water (solvent A) and 90:10 (vol/vol) isopropanol/acetonitrile was performed as described before (61). As previously outlined the LC-electrospray ionization (ESI)-HRMS^2^ analyses were achieved by coupling the LC system to a hybrid quadrupole time-of-flight (QTOF) mass spectrometer Agilent 6538 (Agilent Technologies) equipped with dual electrospray ionization (ESI) (62). For quantifications, 2 μL of internal standards (EquiSPLASH LIPIDOMIX, 330731-1EA) were added prior to extraction. The files generated by Agilent (*.d) were converted to the *.mzML format using MSConvert (63). Subsequently, the software MS-DIAL version 5.1 was employed for data processing and lipid annotation (64). The peak height was utilized as the intensity measure for each annotated lipid in the mass spectra. The nomenclature for lipid sub-classes adheres to the definition provided in (65). Up to nine assays were carried out for lipid analysis during three independents assays. Ouliers analyses was performed via Prism Software V 9.0

### Statistical analysis

MetaboAnalyst 5.0 [**63**] was used to estimate variation across the sample group (PLS-DA and volcano plot). For the Volcano plot, the fold change threshold was 3.0 and the *p*-value threshold was 0.05. Significance was analyzed using ANOVA, and Tukey’s was used as a *post hoc* test. Graphs were made using Prism Software V 9.0. The results were considered significant for a *p* value of ≤0.05.

## Data availability

All data associated with this study are provided in the article.

## ACKNOWLEDGEMENTS

Our gratitude extends to the European Regional Development Fund and the Region of Picardy (CPER 2007– 2020) for their support. We also appreciate UAR 2014—US 41—Plateformes Lilloises en Biologie et Santé and PAGés-P3M for providing the necessary scientific and technical setting that facilitated the completion of this study.

