## Supplementary material for "Unraveling the Interplay of Temperature Adaptation, Lipidomics, and Environmental Factors in *Acinetobacter baumannii* Clinical Strains": Sup1

```
AB5075 1    GTCGTAACAAGGTAGCCGTAGGGGAACCTGCGGCTGGATCACCTCCTTAACGAAAGATTG
ABVal1 1    -----------------------------------------TACCTCCTTACGAAGATTG
ABVal2 1    ---------------------------------------------ACCTTACGACGATTG
ABVal3 1    --------------------------------------------------AGAAAGATTG
ABVal4 1    -----------------------------------------------------ATGATTG
ABVal5 1    ------------------------------------------ACTCCTTACGAAAGATTG

AB5075 61   ACGATTGGTAAGAATCCACAACAAGTTGTTCTTCATAGATGTATCTGAGGGTCTGTAGCT
ABVal1 20   ACGATTGGTAAGAATCCACAACAAGTTGTTCTTCATAGATGTATCTGAGGGTCTGTAGCT
ABVal2 16   ACGATTGGTAAGAATCCACAACAAGTTGTTCTTCATAGATGTATCTGAGGGTCTGTAGCT
ABVal3 11   ACGATTGGTAAGAATCCACAACAAGTTGTTCTTCATAGATGTATCTGAGGGTCTGTAGCT
ABVal4 8    ACGATTGGTAAGAATCCACAACAAGTTGTTCTTCATAGATGTATCTGAGGGTCTGTAGCT
ABVal5 19   ACGATTGGTAAGAATCCACAACAAGTTGTTCTTCATAGATGTATCTGAGGGTCTGTAGCT

AB5075 121  CAGTTGGTTAGAGCACACGCTTGATAAGCGTGGGGTCACAAGTTCAAGTCTTGTCAGACC
ABVal1 80   CAGTTGGTTAGAGCACACGCTTGATAAGCGTGGGGTCACAAGTTCAAGTCTTGTCAGACC
ABVal2 76   CAGTTGGTTAGAGCACACGCTTGATAAGCGTGGGGTCACAAGTTCAAGTCTTGTCAGACC
ABVal3 71   CAGTTGGTTAGAGCACACGCTTGATAAGCGTGGGGTCACAAGTTCAAGTCTTGTCAGACC
ABVal4 68   CAGTTGGTTAGAGCACACGCTTGATAAGCGTGGGGTCACAAGTTCAAGTCTTGTCAGACC
ABVal5 79   CAGTTGGTTAGAGCACACGCTTGATAAGCGTGGGGTCACAAGTTCAAGTCTTGTCAGACC

AB5075 181  CACCATGACTTTGACTGGTTGAAGTTATAGATAAAAGATACATGATTGATGATGTAAGCT
ABVal1 140  CACCATGACTTTGACTGGTTGAAGTTATAGATAAAAGATACATGATTGATGATGTAAGCT
ABVal2 136  CACCATGACTTTGACTGGTTGAAGTTATAGATAAAAGATACATGATTGATGATGTAAGCT
ABVal3 131  CACCATGACTTTGACTGGTTGAAGTTATAGATAAAAGATACATGATTGATGATGTAAGCT
ABVal4 128  CACCATGACTTTGACTGGTTGAAGTTATAGATAAAAGATACATGATTGATGATGTAAGCT
ABVal5 139  CACCATGACTTTGACTGGTTGAAGTTATAGATAAAAGATACATGATTGATGATGTAAGCT

AB5075 241  GGGGACTTAGCTTAGTTGGTAGAGCGCCTGCTTTGCACGCAGGAGGTCAGGAGTTCGACT
ABVal1 200  GGGGACTTAGCTTAGTTGGTAGAGCGCCTGCTTTGCACGCAGGAGGTCAGGAGTTCGACT
ABVal2 196  GGGGACTTAGCTTAGTTGGTAGAGCGCCTGCTTTGCACGCAGGAGGTCAGGAGTTCGACT
ABVal3 191  GGGGACTTAGCTTAGTTGGTAGAGCGCCTGCTTTGCACGCAGGAGGTCAGGAGTTCGACT
ABVal4 188  GGGGACTTAGCTTAGTTGGTAGAGCGCCTGCTTTGCACGCAGGAGGTCAGGAGTTCGACT
ABVal5 199  GGGGACTTAGCTTAGTTGGTAGAGCGCCTGCTTTGCACGCAGGAGGTCAGGAGTTCGACT

AB5075 301  CTCCTAGTCTCCACCAGAACTTAAGATAAGTTCGGATTACAGAAATTAGTAAATAAAGAT
ABVal1 260  CTCCTAGTCTCCACCAGAACTTAAGATAAGTTCGGATTACAGAAATTAGTAAATAAAGAT
ABVal2 256  CTCCTAGTCTCCACCAGAACTTAAGATAAGTTCGGATTACAGAAATTAGTAAATAAAGAT
ABVal3 251  CTCCTAGTCTCCACCAGAACTTAAGATAAGTTCGGATTACAGAAATTAGTAAATAAAGAT
ABVal4 248  CTCCTAGTCTCCACCAGAACTTAAGATAAGTTCGGATTACAGAAATTAGTAAATAAAGAT
ABVal5 259  CTCCTAGTCTCCACCAGAACTTAAGATAAGTTCGGATTACAGAAATTAGTAAATAAAGAT

AB5075 361  TGAGATCTTGGTTTATTAACTTCTGTGATTTCATTATCACGGTAATTAGTGTGATCTGAC
ABVal1 320  TAAGATCTTGGTTTATTAACTTCTGTGATTTCATTATCACGGTAATTAGTGTGATCTGAC
ABVal2 316  TGAGATCTTGGTTTATTAACTTCTGTGATTTCATTATCACGGTAATTAGTGTGATCTGAC
ABVal3 311  TAAGATCTTGGTTTATTAACTTCTGTGATTTCATTATCACGGTAATTAGTGTGATCTGAC
ABVal4 308  TGAGATCTTGGTTTATTAACTTCTGTGATTTCATTATCACGGTAATTAGTGTGATCTGAC
ABVal5 319  TGAGATCTTGGTTTATTAACTTCTGTGATTTCATTATCACGGTAATTAGTGTGATCTGAC

AB5075 421  GAAGACACATTAACTCATTAACAGATTGGCAAAATTGAGTCTGAAATAAATTGTTCACTC
ABVal1 380  GAAGACACATTAACTCATTAACAGATTGGCAAAATTGAGTCTGAAATAAATTGTTCACTC
ABVal2 376  GAAGACACATTAACTCATTAACAGATTGGCAAAATTGAGTCTGAAATAAATTGTTCACTC
ABVal3 371  GAAGACACATTAACTCATTAACAGATTGGCAAAATTGAGTCTGAAATAAATTGTTCACTC
ABVal4 368  GAAGACACATTAACTCATTAACAGATTGGCAAAATTGAGTCTGAAATAAATTGTTCACTC
ABVal5 379  GAAGACACATTAACTCATTAACAGATTGGCAAAATTGAGTCTGAAATAAATTGTTCACTC

AB5075 481  AAGAGTTTAGGTTAAGCAATTAATCTAGATGAATTGAGAACTAGCAAATTAACTGAATCA
ABVal1 440  AAGAGTTTAGGTTAAGCAATTAATCTAGATGAATTGAGAACTAGCAAATTAACTGAATCA
```

```
ABVal2 436 AAGAGTTTAGGTTAAGCAATTAATCTAGATGAATTGAGAACTAGCAAATTAACTGAATCA
ABVal3 431 AAGAGTTTAGGTTAAGCAATTAATCTAGATGAATTGAGAACTAGCAAATTAACTGAATCA
ABVal4 428 AAGAGTTTAGGTTAAGCAATTAATCTAGATGAATTGAGAACTAGCAAATTAACTGAATCA
ABVal5 439 AAGAGTTTAGGTTAAGCAATTAATCTAGATGAATTGAGAACTAGCAAATTAACTGAATCA

AB5075 541 AGCGTTTTGGTATGTGAATTTAGATTGAAGCTGTACGGTGCTTAAGTGCACAGTGCTCTA
ABVal1 500 AGCGTTTTGGTATGTGAATTTAGATTGAAGCTGTACGGTGCTTAAGTGCACAGTGCTCTA
ABVal2 496 AGCGTTTTGGTATGTGAATTTAGATTGAAGCTGTACGGTGCTTAAGTGCACAGTGCTCTA
ABVal3 491 AGCGTTTTGGTATGTGAATTTAGATTGAAGCTGTACGGTGCTTAAGTGCACAGTGCTCTA
ABVal4 488 AGCGTTTTGGTATGTGAATTTAGATTGAAGCTGTACGGTGCTTAAGTGCACAGTGCTCTA
ABVal5 499 AGCGTTTTGGTATGTGAATTTAGATTGAAGCTGTACGGTGCTTAAGTGCACAGTGCTCTA

AB5075 601 AACTGAAATGTTGAAGTTACTAACTTGTAGGTAACATCGACTGTTTGGGGTTGTATAGTC
ABVal1 560 AACTGAAATGTTGAAGTTACTAACTTGTAGGTAACATCGACTGTTTGGGGTTGTATAGTC
ABVal2 556 AACTGAAATGTTGAAGTTACTAACTTGTAGGTAACATCGACTGTTTGGGGTTGTATAGTC
ABVal3 551 AACTGAAATGTTGAAGTTACTAACTTGTAGGTAACATCGACTGTTTGGGGTTGTATAGTC
ABVal4 548 AACTGAAATGTTGAAGTTACTAACTTGTAGGTAACATCGACTGTTTGGGGTTGTATAGTC
ABVal5 559 AACTGAAATGTTGAAGTTACTAACTTGTAGGTAACATCGACTGTTTGGGGTTGTATAGTC

AB5075 661 AAGTAATTAAGTGCATGTGGTGGATGCCTTGGCAGTCAGAGGCGATGAAAGACGTGATAG
ABVal1 620 AAGTAATTAAGTGCATGTGGTGGATGCCTTGGCAGTCAGAGGCGATGAAAGACGTGATAG
ABVal2 616 AAGTAATTAAGTGCATGTGGTGGATGCCTTGGCAGTCAGAGGCGATGAAAGACGTGATAG
ABVal3 611 AAGTAATTAAGTGCATGTGGTGGATGCCTTGGCAGTCAGAGGCGATGAAAGACGTGATAG
ABVal4 608 AAGTAATTAAGTGCATGTGGTGGATGCCTTGGCAGTCAGAGGCGATGAAAGACGTGATAG
ABVal5 619 AAGTAATTAAGTGCATGTGGTGGATGCCTTGGCAGTCAGAGGCGATGAAAGACGTGATAG

AB5075 721 CCTGCGAAAAGCTCCGGGGAGGCGGCAAATATCCTTTGATCCGGAGATGTCTGAATGGGG
ABVal1 680 CCTGCGAAAAGCTCCGGGGAGGCGGCAAATATCCTTTGATCCGGAGATGTCTGAAC----
ABVal2 676 CCTGCGAAAAGCTCCGGGGAGGCGGCAAATATCCTTTGATCCGGAGATGTCTGATG----
ABVal3 671 CCTGCGAAAAGCTCCGGGGAGGCGGCAAATATCCTTTGATCCGGAGATGTCTGATGGGG-
ABVal4 668 CCTGCGAAAAGCTCCGGGGAGGCGGCAAATATCCTTTGATCCGGAGATGTCTGATGG---
ABVal5 679 CCTGCGAAAAGCTCCGGGGAGGCGGCAAATATCCTTTGATCCGGAGATGTCTGATGGGGG

AB5075 781 GAACCC-
ABVal1 735 -------
ABVal2 731 -------
ABVal3 729 -------
ABVal4 724 -------
ABVal5 739 GAACCCC
```
